# Multi-character assessment of caryopsis in diverse common wheat germplasm, including high protein breeds

**DOI:** 10.1101/2024.10.26.620432

**Authors:** Romuald Kosina, Paulina Tomaszewska

## Abstract

In 15 varieties and hybrids of common wheat, including high-protein, varieties such as Atlas 66, Nap Hal, and Lancota, the microstructural characters of the caryopsis were analysed. Among the wheats, a high variability in the correlation matrices of characters was observed. A variable relationship between the specific gravity of the caryopsis and the size of the endosperm cavity was demonstrated. Some principal components (I, II, IV and V) contained the majority of the information regarding the weight, quality, vitreousness, and aleurone layer of the caryopsis, respectively. In the ordination space of nonmetric multidimensional scaling (nmMDS), two groups of wheat were distinguished: high-protein and low-protein subaleurone layers. Endosperm morphotypes were characterised by the proportion of two types of starch tissue, cylindric and isodiametric. In the nmMDS ordination space, wheat was described by the frequency distributions of six endosperm cavity morphotypes, with more (leptokurtosis) or less (platykurtosis) cumulated variation. In the case of endosperm developmental anomalies, the intrusive growth of cell walls, particularly in the aleurone layer, was identified as playing a significant role in the formation of endosperm mosaic architecture and domains. Varietal developmental specificity was discovered in the development of acellular starch domains within the endosperm tissue and starch grain storage in the cavity (as observed in cv. Purdue).

## 1. Introduction

Wheat, alongside maize and rice, is one of the world’s primary cereals for human nutrition [1]. The production status of these cereals globally remains stable, with a trend of slow yield growth [2]. In the temperate climate zone, wheat is the dominant cereal. A crucial aspect of its nutritional value is the protein content and the presence of essential amino acids in the grain. During the 1970s and 1980s, a global research project conducted at the University of Nebraska, Lincoln, utilised high-protein varieties such as Atlas 66 and Nap Hal as sources of genes to develop new lines with enhanced nutritional value [3]. These varieties continue to be regarded as valuable materials in wheat breeding, particularly in light of the challenges associated with improving food quality [1,4]. This justifies the detailed investigation of the structure and quality of common wheat grains, with particular emphasis on the relationships between different caryopsis tissues and their influence on the accumulation of assimilates in the form of starch and/or protein. A comprehensive synthesis of the structure and quality of the wheat caryopsis was provided by [5]. The caryopsis is an organ where various components influence its final form. Its complete description was obtained following the analysis of many characters, including both structural and non-structural data. Multi-variate analyses of the caryopsis have been conducted previously for various wheats [6,7]. Multi-variate analysis enables the synthesis of data and the taxonomic description of different wheat breeding units, such as pure varieties and hybrids. The covariation of caryopsis characters related to morphology and quality can be represented by principal components, which serve as synthetic markers of characters in wheat breeding units [6]. The differentiation of caryopsis tissues depends on their position relative to the shoot axis and has been studied in its early stages by [8–11]. Important developmental events include the morphogenesis of the ventral side near the transfer tissue complex, the dorsal side of the caryopsis, the central part and the two lateral lobes. The tissues in the starchy endosperm differ in the morphogenesis of the cylindrical and isodiametric cells that compose them. The final stage in the ‘filling’ of the caryopsis involves the closure or non-closure of the space between the dorsal and ventral areas [11]. The morphogenesis of the endosperm cavity, starchy endosperm, and transfer system of the caryopsis reflects the developmental interactions among these structures [12,13]. Developmental events in the caryopsis should be understood in the context of the domain nature of the endosperm, which is revealed after the elimination of defective nuclei in the embryo sac syncytium, particularly in hybrid forms. This demonstrates the exceptional developmental and morphological complexity of the caryopsis [14–16]. The final shape of the caryopsis is influenced by the morphogenesis of the chalazal transfer area and the adjacent endosperm cavity. The caryopsis forms between leafy glumellae, which create a ‘floral cavity’ that limits its growth [17]. Among wheat species, it has been established that the sclerification and adherence of the glumellae to the caryopsis significantly affect the shape and size of the endosperm cavity [18]. Seed size is also influenced by the stiffening of the testa [19], which in wheat may be related to the maturation of fruit tissues in interaction with the glumellae forming the flower cavity. The apoptotic degradation of maternal tissues, including the nucellus [20], has been considered a source of additional substrates supplied to the endosperm through the endosperm cavity, which is part of the caryopsis apoplast [21,22].

The developmental variations of the wheat caryopsis guided the objective of this study towards a multivariate analysis of caryopsis microstructure in wheat varieties and hybrids, with a focus on cell and tissue differentiation as well as the covariation of structure and nutritional quality.

## 2. Materials and methods

### 2.1. Materials

The study was conducted on 15 accessions (cultivars and hybrids) of *Triticum aestivum* L. cultivated at the Plant Breeding Station Pustków Żurawski on loess soil. All accessions were planted under uniform soil and climatic conditions, each on a single small plot. Winter and intermediate forms were sown in autumn, while spring forms were planted in the spring. The material was treated as a design for a completely randomised one-way classification. The intraspecific units and their symbols, used in the text and diagrams, are listed in Table 1. Data on the breeding origin were cited according to [23] and were also provided by V.A. Johnson (pers. comm.) (Table 1).

**Table 1.** Cultivars and hybrids of common wheat.

| Cultivars | Symbol | Breeding origin | Donor |
| --- | --- | --- | --- |
| Atlas 66 | A | Frondoso//Redhart 3/Noll 28 | Lincoln, USA |
| C.I. 13449 | CI | Norin 10/Brevor, F<sub>5</sub>, a sib line to Brevor | Pullman, USA |
| Lancota | L | Atlas 66/Comanche//Lancer | " |
| Nap Hal | NH | Indian landrace | India |
| Purdue 4930A6-28-2-1 | P | A selection line from Frondoso | Oklahoma, USA |
| Octavia | O | Stupicka Vouska/Koga | Czechoslovakia |
| Ostka Popularna | OP | Ostka Kleszczewska/?/Heines Kolben /?/<br>von Rumkers Frühreifer Dickkopf | Poland |
| Yuma experiment numbers for hybrids (H) | Symbol | Breeding origin | Donor |
| 11847 | H1 | Nap Hal/Justin | Lincoln, USA |
| 11507 | H2 | Nap Hal/CIMMYT 8156 | " |
| 10699 | H3 | Nap Hal/C.I. 13449, F<sub>6</sub> | " |
| 10100 | H4 | Nap Hal/C.I. 13449, F<sub>6</sub> | " |
| 21061 | H5 | C.I. 13449/Centurk | " |
| 10121 | H6 | Nap Hal/Atlas 66, F<sub>7</sub> | " |
| 10841 | H7 | Nap Hal/C.I. 13449, F<sub>6</sub> | " |
| 11574 | H8 | Nap Hal/CB 113 | " |

Data on breeding origin are given according to [23]Zeven and Zeven (1976) and provided by V.A. Johnson (personal comm.) from experiments at Yuma, University of Nebraska, Lincoln

### 2.2. Evaluation of characters

For each wheat accession, a random sample of ripe caryopses (*n* = 100) was evaluated based on the following nine characters (character symbols are given in brackets):

1. Weight of caryopsis (W, mg).
2. Specific gravity (density) (SG, g/cm^3^) This was calculated using the following formula: ρ_caryopsis_ (SG) = *W*_a_/*V*_caryopsis_, with the data derived from the volume of the caryopsis determined from the mass of the caryopsis in atmospheric air (*W*_a_) and in absolute alcohol (*W*_etoh_), with a specific gravity of ρ_etoh_ = 0.7892. The formula used was *W*_a_ - *W*_etoh_/ρ_etoh_ = *V*_caryopsis_.
3. Type of endosperm (MV); five types of mealy : vitreous endosperm percentage noted on a transverse fracture of the caryopsis: **1** (mealy) – 100% : 0%, **2** – 75% : 25%, **3** – 50% : 50%, **4** – 25% : 75%, **5** (vitreous) – 0% : 100%.
4. Width of the crease (WC, mm).
5. Thickness of aleurone layer (TA, µm).
6. Thickness of lateral high protein (HP) subaleurone layer (TLS, µm).
7. Thickness of dorsal high protein (HP) subaleurone layer (TDS, µm).
8. Surface area of endosperm cavity (SAC, cm^2^). The cavity was drawn using an Amplival polarising microscope with the MNR1 drawing microscope ocular (PZO Warsaw), and its surface area was measured on the drawing using a PL1 polar planimeter (PZO Warsaw).
9. Width of endosperm cavity (WEC, µm).

Characters 1 and 2 were measured on the whole mature caryopsis, character 3 on its fracture, and characters 4 - 9 on cross-sections of the central part of the caryopsis, which was fixed in FAA fixative (formalin : acetic acid : ethanol) and cut on a microtome using a freezing stage with a TOS-11 selenium rectifier (V/O Medèksport, Moskva). To screen the total protein content, cross-sections were stained with a 1% aqueous solution of bromophenol blue, following the method of [24], washed in tap water, and mounted in glycerine on semi-permanent slides. Protein morphotypes, ranging from low protein (LP) to high protein (HP) subaleurone layers, were recorded. Measurements of the characters and photographic documentation were made using an Amplival polarising microscope (Carl Zeiss, Jena), as well as Olympus BX-50 and Zeiss Axio Imager M2 epifluorescence microscopes. Additionally, the variability of the endosperm cavity shape was described by six morphotypes (A, B, C, D, E, F).

### 2.3. Numerical taxonomy

For the research data, correlation matrices were created for each accession, and principal components analysis (PCA) was performed for nine wheat breeds using the character sets, following the method of [25]. For the wheat accessions (operational taxonomic units, OTUs) and their characters, matrices of average taxonomic distances (Euclidean distances) were calculated according to the Q and R techniques [26], using programs provided by [27]. These matrices were used as input data for non-metric multidimensional scaling (nmMDS) [28]. The nmMDS symmetric matrix of distances in the final configuration space was then used to calculate edges in a minimum-length spanning tree (MST). The tree was plotted, as is commonly practiced, onto a 3D-sphere according to the proposal of [26].

## 3. Results

### 3.1. Characteristics of caryopsis in common wheat

#### 3.1.1. Variation of character correlations

Two matrices of Pearson’s significant correlation coefficients between the studied caryopsis characters reveal a wide range of values - seven significant correlations in the H7 hybrid and 26 in the Lancota (L) variety (Table 2). The significant coefficients in the H7 matrix also appear in the L matrix with similar values. In the H7 matrix, the highest correlation significance is observed between the characters W-SAC and TLS-TDS, whereas, in the L matrix, the covariation of many characters is strongly marked, showing high levels of correlation significance. Other OTUs present intermediate numbers of significant coefficients. The number of significant coefficients across both varieties and hybrids is similar, indicating a comparable degree of covariation of characters. For instance, the H7 hybrid and the Atlas 66 (A) variety show similarly low numbers of significant coefficients, with seven and nine, respectively. The correlation coefficients between caryopsis weight (W) and the characters SG, TA, TLS, and TDS suggest that larger caryopses tend to have lower specific gravity and thicker aleurone and HP subaleurone layers. A comparison of the most significant correlation coefficients in the Lancota variety for the characters W-WC, W-TLS, W-TDS, W-SAC, W- WEC, SG-MV, and WC-WEC with corresponding coefficients in other OTUs shows that some of these coefficients are statistically insignificant in many OTUs. Of particular interest are the correlations involving the SG character (specific gravity) and other caryopsis traits. Significant positive correlations were found between SG and the MV (mealy/vitreous endosperm) character in many OTUs, while negative correlations were noted between SG and WC (width of crease) and WEC (width of endosperm cavity). SG’s correlations with TA, TLS, TDS, and SAC exhibit a dual nature (+ or −), indicating different covariation patterns between these structures and the other components of the caryopsis (such as maternal tissues, starchy endosperm, and the embryo), all of which contribute to the specific gravity of the caryopsis. An analysis of the significant coefficients for the SG-SAC relationship reveals that in OTUs with large endosperm cavities, the SG value decreases (negative correlation), while in OTUs with small cavities, the SG value increases (positive correlation). The range of SAC mean values across OTUs varies from 3 (OP, NH, H4) to 11 (L). This variable relationship indicates two developmental trends of the cavity: one related to its growth and metabolite content, and the other influencing specific gravity. Positive W-SAC correlations indicate that as caryopsis weight increases, so does the volume of the endosperm cavity; these correlation coefficients show extreme values, from *r* = 0.20 in H2 to *r* = 0.71 in CI. In 33%of wheat breeds, this correlation is insignificant.

**Table 2.** Two exemplary matrices of significant coefficients of Pearson’s correlation with extreme their numbers (7 and 26) in H7 hybrid and Lancota variety (*n* = 100)

| Hybrid H7<br>Characters | W | SG | MV | WC | TA | TLS | TDS | SAC | WEC |
| --- | --- | --- | --- | --- | --- | --- | --- | --- | --- |
| W | 1.00 |  |  |  |  |  |  |  |  |
| SG | -.22* | 1.00 |  |  |  |  |  |  |  |
| MV | ns | .20* | 1.00 |  |  |  |  |  |  |
| WC | ns | ns | ns | 1.00 |  |  |  |  |  |
| TA | .31** | ns | ns | ns | 1.00 |  |  |  |  |
| TLS | ns | ns | ns | ns | ns | 1.00 |  |  |  |
| TDS | ns | ns | ns | ns | ns | .48*** | 1.00 |  |  |
| SAC | .38*** | ns | ns | ns | ns | ns | ns | 1.00 |  |
| WEC | ns | ns | ns | .79*** | ns | ns | ns | .22* | 1.00 |
| Lancota (L)<br>Characters | W | SG | MV | WC | TA | TLS | TDS | SAC | WEC |
| W | 1.00 |  |  |  |  |  |  |  |  |
| SG | -.23* | 1.00 |  |  |  |  |  |  |  |
| MV | .24* | .55*** | 1.00 |  |  |  |  |  |  |
| WC | .69*** | -.31** | ns | 1.00 |  |  |  |  |  |
| TA | .24* | ns | ns | ns | 1.00 |  |  |  |  |
| TLS | .49*** | ns | .44*** | .42*** | ns | 1.00 |  |  |  |
| TDS | .50*** | ns | .31** | .24* | .30** | .47*** | 1.00 |  |  |
| SAC | .47*** | -.29** | ns | .39*** | .25* | .29** | .23* | 1.00 |  |
| WEC | .72*** | ns | .22* | .72*** | ns | .46*** | .35*** | .29** | 1.00 |

\*, \*\*, \*\*\* - significance levels of correlation coefficients at $\alpha = 0.05, 0.01, 0.001$ , respectively ns – non-significant coefficients

#### 3.1.2. Character loadings in principal components

To illustrate the level of information on the variation of characters described by the principal components (loadings), a threshold of 40% and above was assumed (Table 3). The 50% threshold was abandoned because, at that level, the information on the characters of the Purdue variety was not fully represented within the principal components. The PCA analysis was conducted for nine OTUs. The following components exhausted information about different sets of characters as follows:

- Component I - W, WC, WEC, and also SG (a component of caryopsis weight, width of crease, and width of endosperm cavity).
- Component II - TLS, TDS (a component of grain quality).
- Component III - Information about various characters.
- Component IV - MV (component of mealy/vitreous endosperm information).
- Component V - TA (component of aleurone layer thickness).

**Table 3.** Hotelling’s principal components analysis for the caryopsis characters of nine breeds of common wheat.

| Breeds of<br>common wheat | C o m p o n e n t s |  |  |  |  |  | Sum of<br>variation<br>(%) | Sum of<br>variation<br>(%) |
| --- | --- | --- | --- | --- | --- | --- | --- | --- |
|  | I | % * | II | III | IV | V | I-V | I-IX |
| A | WC-WEC | 26 | TLS-TDS | - | MV | TA | 77 | 86 |
| CI | W-SAC | 31 | WC-WEC | SG-MV | - | SG | 84 | 91 |
| L | W-WC-TLS-WEC | 40 | SG-MV | TA | SAC | - | 87 | 92 |
| NH | W-SG-WC-WEC | 33 | TLS | SAC | MV | TA | 83 | 90 |
| P | W-TLS-TDS | 28 | MV | WC |  | TA | 81 | 88 |
| O | W | 29 | WC-WEC | TDS | MV | - | 82 | 89 |
| OP | W-WC-WEC | 28 | SG | TDS | MV | - | 84 | 91 |
| H2 | W-SG-WC-WEC | 34 | TDS | SAC | MV | - | 82 | 89 |
| H3 | W-WC-WEC | 33 | TLS-TDS | TA | - | - | 85 | 91 |

I-IX – principal components; \* - % of variation of nine characters contained in the first component:

Components I, II, IV, and V provided the most consistent information regarding the variation of characters across the set of wheat studied. The wheat accessions distinctly differ in the sets of characters represented by these five components. In particular, the Atlas 66 (A) variety exhibits the lowest level of information coverage, with 77% of the total variation being exhausted. The extreme difference in the level of information exhausted for the nine characters by Component I is noticeable between the Atlas 66 (26%) and Lancota (40%) varieties.

#### 3.1.3. Distribution of caryopsis characters in nmMDS ordination space

Two groups of characters, initially described by the matrix of average taxonomic distances, can be identified in the minimum spanning tree diagram (Fig. 1). The first group consists of the characters W, TA, TDS, and TLS, which are notably dispersed across the diagram. The last three characters, due to their origin from tangential divisions and their proximity within the endosperm, are undoubtedly developmentally correlated. The second group includes the remaining caryopsis characters, which are strongly clustered and describe features such as the caryopsis crease and endosperm cavity (WC, SAC, WEC), specific gravity (SG), and the proportion of mealy versus vitreous endosperm (MV). These characters are described by the maximum values on the *x*-axis and the minimum values on the *y*- and *z*-axis in the ordination space. The spatial relationships of these characters are partially consistent with the information on their variation contained in the principal components.

**Fig. 1.**
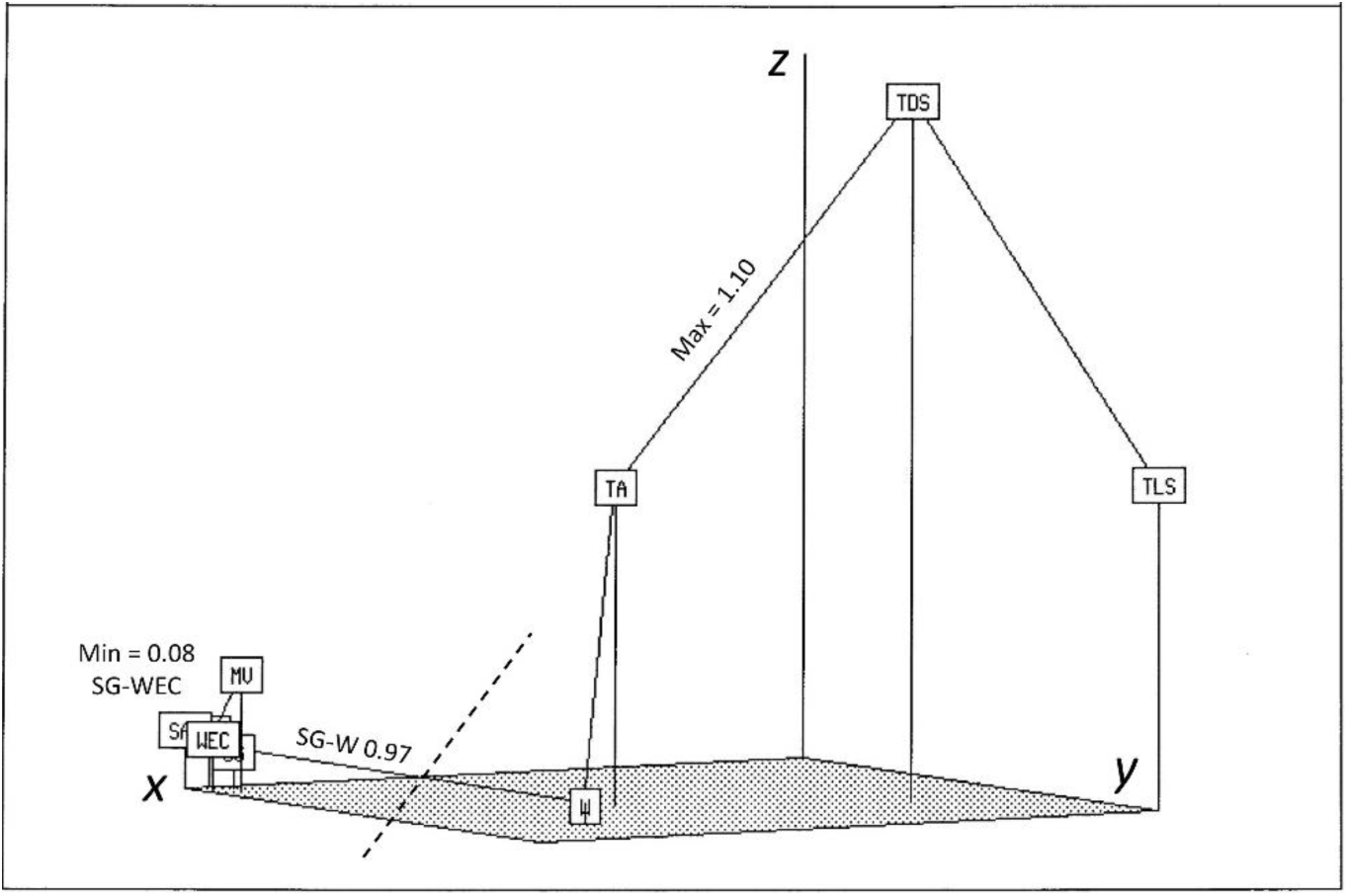
Minimum spanning tree of nine caryopsis characters in an ordination space defined by ***x***-, ***y***-, and ***z***-axes, and created by application of Kruskal’s non-metric multidimensional scaling method [27](Rohlf 1994). Characters were described by 15 arithmetic averages of common wheat breeds. In the tree minimum and maximum distances are shown (see Materials and methods for labels of characters).

#### 3.1.4. Distribution of common wheat OTUs in nmMDS ordination space

In the ordination space, OTUs A and H3, which belong to the cluster characterised by the thickest HP-subaleurone layer in both the dorsal and lateral positions are extreme to the distant O variety with a thin LP-subaleurone layer (Fig. 2). Other OTUs display an intermediate status regarding these subaleurone endosperm characteristics, including a cluster of three varieties: CI, OP, and L. The ‘HP-subaleurone’ cluster, which also contains several hybrids (H1–H5) bred specifically to increase protein content, demonstrates breeding progress as reflected in the changes in the wheat caryopsis microstructure.

**Fig. 2.**
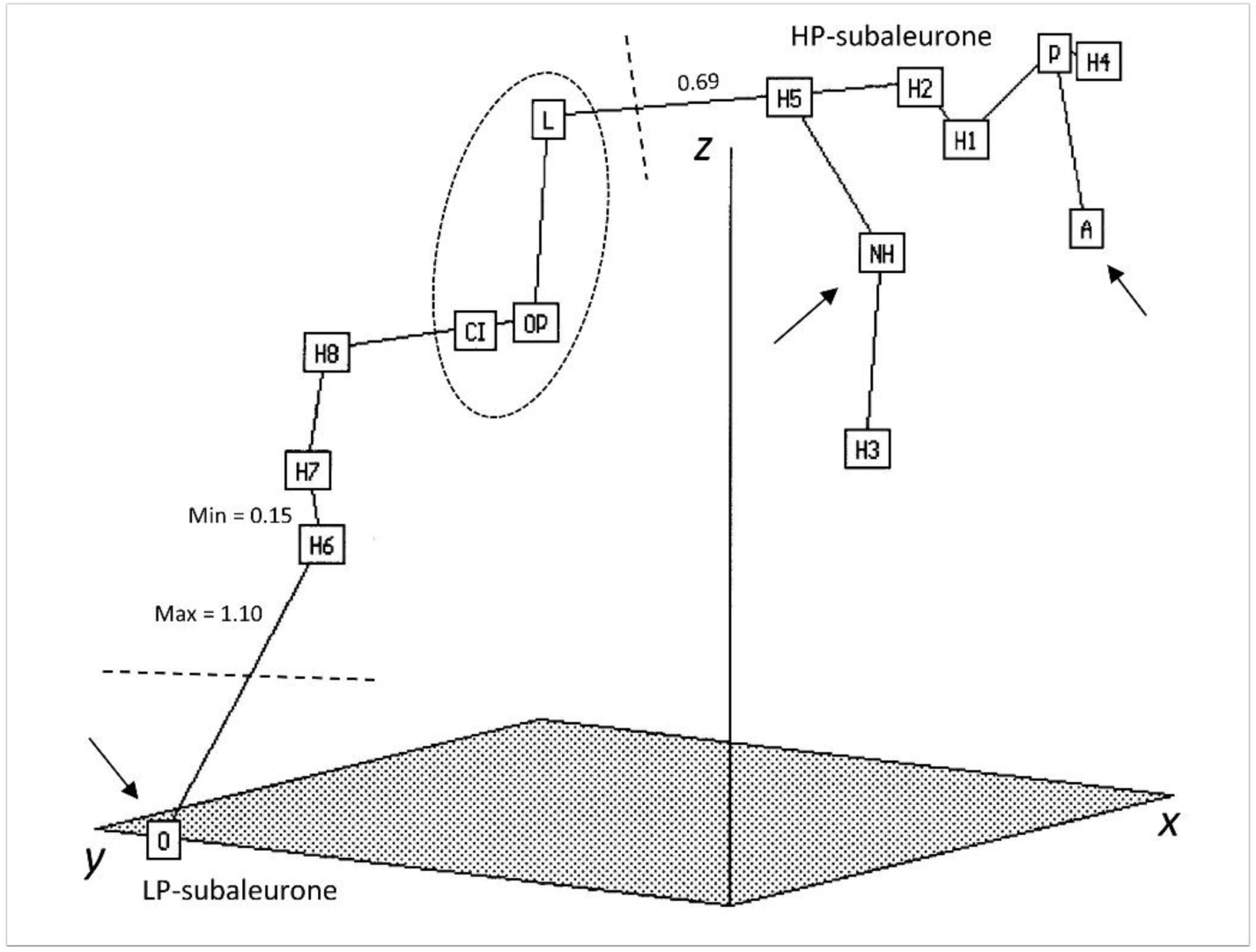
Minimum spanning tree of 15 OTUs of common wheats described by arithmetic averages of nine characters of caryopsis in an ordination space defined by *x*-, *y*-, and *z*-axes, and created by the application of Kruskal’s non-metric multidimensional scaling method [27](Rohlf 1994). In the tree, minimum and maximum distances are presented. Labels for OTUs in M&M.

### 3.2. Characteristics of caryopsis anatomy and its irregularities

#### 3.2.1 Characteristics of the endosperm structure

The relationships between filial cells and tissues enclosed within the maternal body (pericarpium) are crucial for the quality assessment of wheat caryopsis and are partially depicted in Fig. 3. In the transverse section of the caryopsis, the spatial organization of cylindrical cells in the dorsal part of the endosperm varies according to the shape of the adjacent endosperm cavity (Fig. 3a, b). In one case, vertical cylindrical cells arranged almost parallel create a horizontal strip of dorsal endosperm adjacent to a flat cavity (*planar cavity*) and a gently convex, flat, or even slightly concave caryopsis ridge. The second arrangement of cylindrical cells resembles the elements of sharp Gothic arches, with the endosperm adjoining a strongly convex or even-keeled caryopsis ridge and a cavity shaped like a Gothic window (*gothic cavity*). These two architectural types of endosperm represent extreme forms, with the wheat grains examined showing a spectrum of variability. The cylindrical endosperm typically transforms in the lateral parts of the caryopsis into endosperm with isodiametric cells (Fig. 3c). However, the distinction between these two types of cells can be challenging due to the lack of clear cell wall boundaries, making it difficult to identify the architectural differences in the endosperm. In genetically stabilised wheat varieties, the starchy endosperm contacts a structurally uniform aleurone layer (Fig. 3d).During the final stage of tangential cytokineses, a single-celled or multi-celled subaleurone layer forms beneath the aleurone layer, composed of short cells with limited elongation growth. In forms with uniform starch synthesis dynamics throughout the starchy endosperm, such as the H3 hybrid (Fig. 3d), this layer is poorly distinguished optically. However, when stained with protein dyes, it is morphologically distinct in high-protein breeds, such as the H4 hybrid (Fig. 3e). The high-protein subaleurone layer (HP- subaleurone) can be continuous with varying thickness, as indicated by positive TLS-TDS correlations. This layer always terminates inthe ventral part of the endosperm cavity, near the transfer system formed by phloem and xylem bundles, pigment strand, and the nucellar projection (Fig. 3f).

**Fig. 3.**
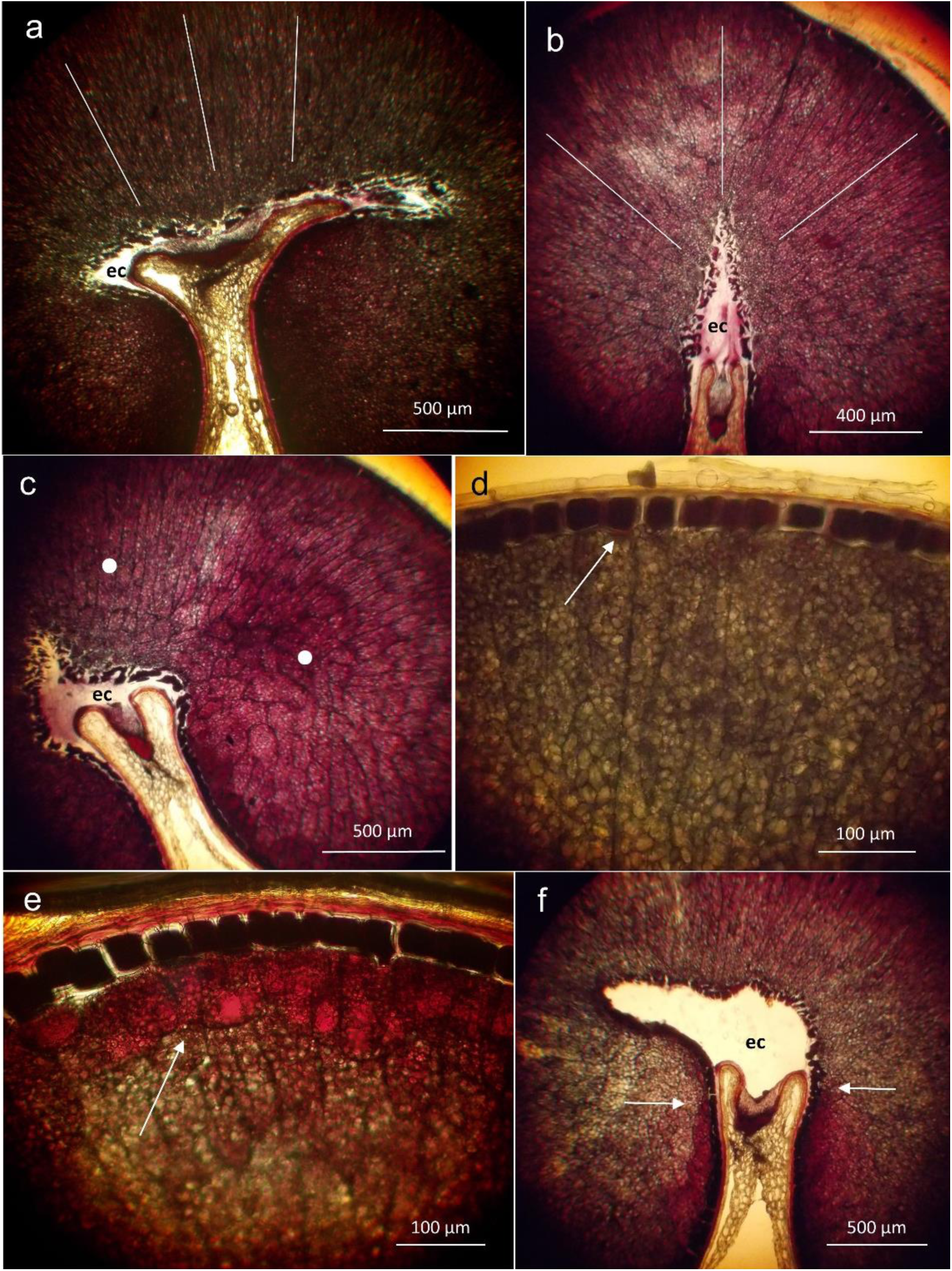
Structural data of common wheat endosperm. **a** – OP, solid lines show the arrangement of cells in the cylindrical endosperm; **b** – A, solid lines as in **a**; **c** – A, two morphotypes of starchy endosperm, cylindric and isodiametric, shown by two white dots; **d** – H3, regular aleurone layer (arrow) and a lack of HP-subaleurone layer; **e** – H4, thick and continuous HP-subaleurone layer (arrow); **f** – L, the ends of the HP-subaleurone layer under the large endosperm cavity (arrows). For **a** – **f** - cross-sections were stained with bromophenol blue.

#### 3.2.2. Characteristics of aleurone–subaleurone area

Two types of starch grains are synthesised in the wheat endosperm: the larger A type and the smaller B type. Regardless of the starch grain type, they are always situated within the protein matrix of the endosperm cells. In the subaleurone layer, B-type grains are predominant, while both A and B grains are found in the centripetal parts of the endosperm. After diagonal cell division, aleurone cells frequently grow intrusively into the subaleurone layer (Fig. 4a).

**Fig. 4.**
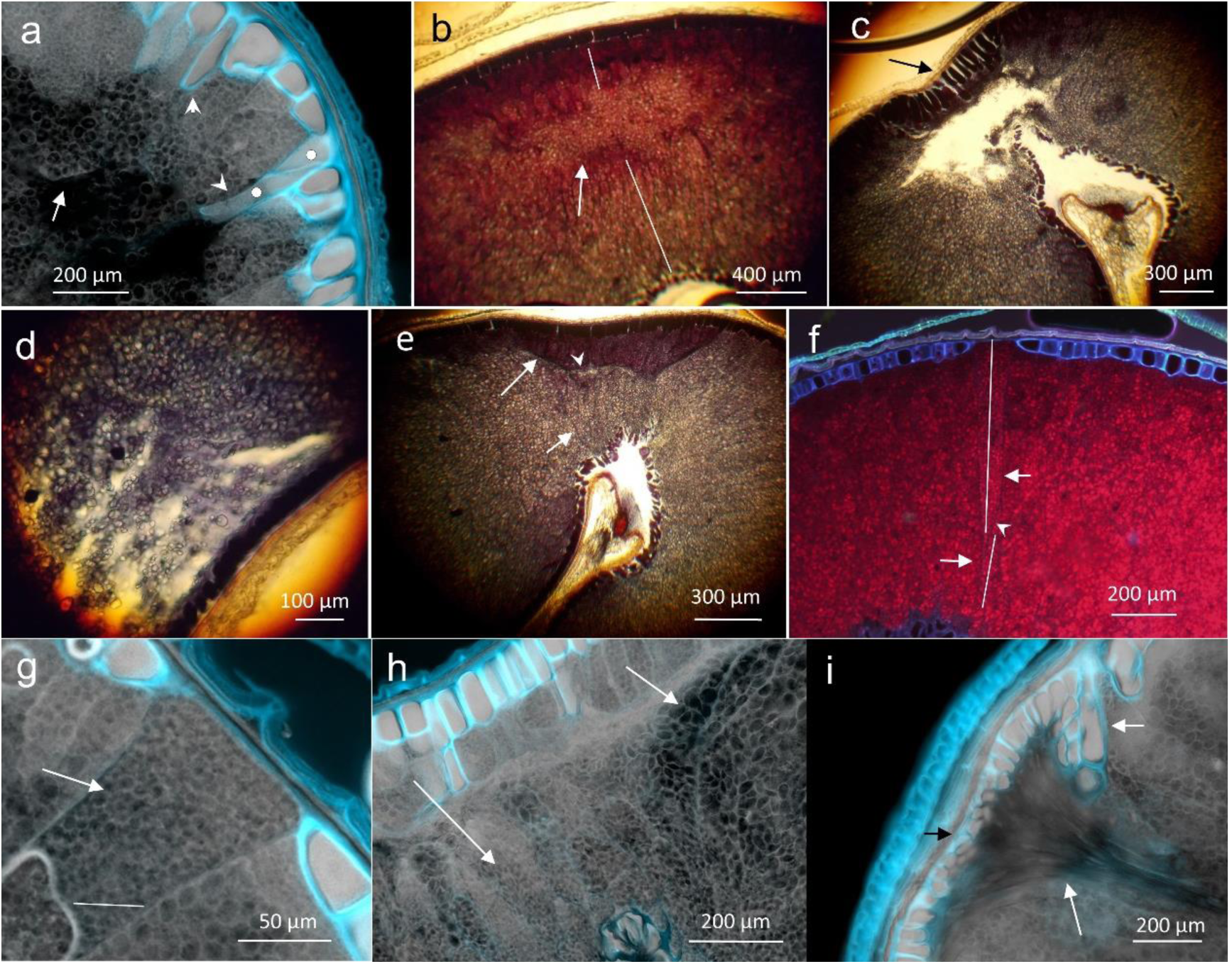
Microstructures in the aleurone-subaleurone area. **a** – H3, starch grains embedded in protein matrix (arrow), intrusive ingrowths in starchy endosperm (arrowheads), two aleurone cells divided by a diagonal division (white dots); **b** – L, an acellular starchy domain (arrow) unequally dividing dorsal-ventral endosperm (white lines); **c** – NH, an anomalous aleurone layer with elongated cells (arrow) neighbouring an empty space in starchy endosperm; **d** – H5, a loose starchy endosperm under dark aleurone layer; **e** – NH, dorsal endosperm composed of two domains (arrows): higher dark HP- subaleurone layer and lower lighter with small amount of protein; both domains are separated by a thin acellular domain (small white dot); **f** – H3, an anomalous two-celled cylindric starchy complex (white lines along cells and arrows) formed within aleurone layer and multiplied by diagonal cytokinesis (arrowhead); **g** – H3, an upper part of the cell complex from **f** showing uniformity of starch grains (arrow), a white line shows an air artefact under cover slip; **h** – H8, two parts of subaleurone starchy endosperm with different starch grains (arrows); **i** – H1, an anomalous domain composed of narrow cells with strong intrusive growth, similar to a bundle of fibers (long arrow). The domain was developed near anomalous aleurone layer (short arrow). Small, very rare starch grains were synthesised in its cells. Autofluorescence of tissues in **a**, **f**, **g**, **h**, **i**, and bromophenol blue staining in others. –

In the studied forms of common wheat, the development of acellular starch domains was observed, typically occurring under the subaleurone layer in the dorsal or dorsal-lateral part. Starch is likely synthesised earlier in these regions, preventing cytokineses within the domain (Fig. 4b). This suggests that energy resources (assimilates) are diverted from endosperm cellularisation towards starch synthesis. The acellular starch domain acts as a separator between the cylindrical endosperm layers - dorsal and ventral - of unequal thickness. The occurrence frequency of this morphotype indicates that it is a rare varietal trait, present in 32% of the Purdue cultivar and 9% of the Lancota variety.

A defect in starch synthesis was also noted as part of a developmental anomaly, characterised by a space in the endosperm, which is filled by the anomalous anticlinal elongation growth of adjacent aleurone cells (Fig. 4c) or bordered by a normal aleurone layer (Fig. 4d). In both cases, these areas lacked cell walls and were similar to acellular domains (Fig. 4b vs. 4c, d), although the latter showed poor starch synthesis.The high-protein subaleurone layer contains small B-type starch grains, which are synthesised in the final stage of endosperm development. This high-protein subaleurone cell morphotype typically develops as either a continuous periclinal stripe or as distinct domains (Fig. 4e). The dorsal HP-domain (Fig. 4e) is developmentally connected to the ventral LP-domain, and both domains are separated by a very narrow acellular layer. The establishment of the starchy endosperm cell type from the onset of endosperm development leads to its presence in the aleurone layer (Fig. 4f), followed by strong elongation and intrusive growth after one diagonal division. Two daughter cells stretch almost the entire dorsal-ventral section and are filled with a single type of starch grains (Fig. 4f, g), exemplifying a one-to-two-celled starch domain.

In areas with developmental anomalies in aleurone cells (Fig. 4a, h), subaleurone cells containing B grains may extend into the endosperm or be lined with cells containing A grains (Fig. 4h). A grains also appear directly under the aleurone layer (Fig. 4a). These examples highlight the mosaic nature of the starchy structure of the endosperm. The unique developmental domain in the H1 hybrid is shown in Fig. 4i. In this case, an area with an aleurone layer developmental anomaly is bordered by a clinal domain of narrow cells, formed after significant elongation and intrusive growth, resembling the growth of sclerenchyma fibres in other plant tissues. These cells contain few starch grains of microscopic dimensions, as the entire energy potential of the domain was utilised for cell division and growth.

#### 3.2.3. Anomalies in the aleurone layer development

In wheat, the developmental rule is the formation of a regular, unicellular aleurone layer. Figure 5 illustrates examples of irregular development in the aleurone layer and its interaction with the starchy endosperm. Anticlinal and periclinal divisions within the aleurone cells represent the final stages of cytokinesis after the periclinal divisions that separate the starchy subaleurone cells (Fig. 5a). Depending on the initial size of the aleurone cell and the extent of tangential growth, clones of small or large cells may form, as well as clones that undergo additional periclinal divisions in the already differentiated aleurone layer (Fig. 5a). In the dorsal part of the Lancota (L) variety, the aleurone layer is absent, and this region is filled by large starch cells (Fig. 5b). In Nap Hal (NH), the original architecture of the caryopsis, characterised by a concave ridge, leads to the development of elongated aleurone cells arranged in the form of an inverted fan in this area (Fig. 5c). This is likely a result of mechanical interactions between tissues during development.

**Fig. 5.**
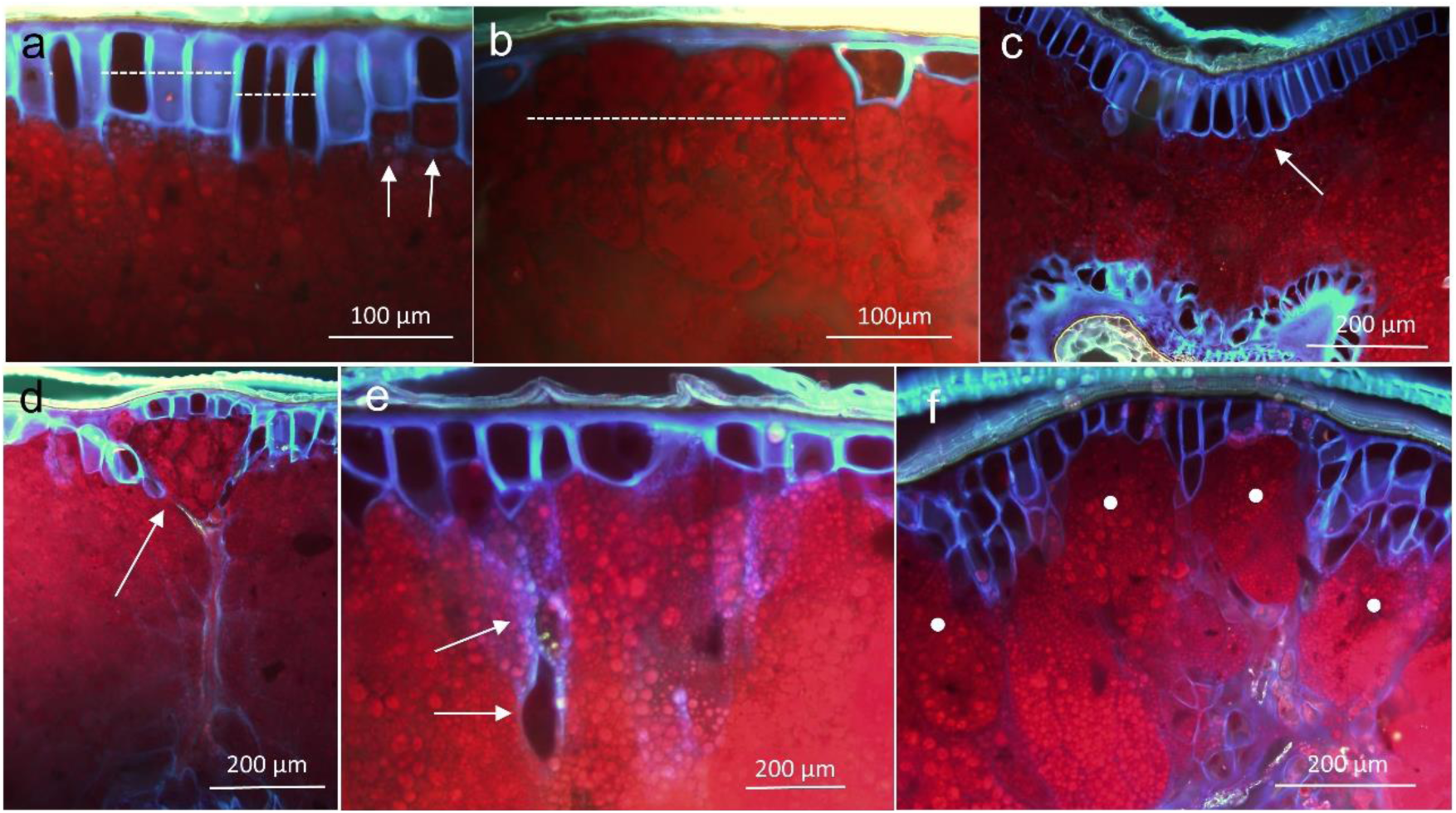
Examples of the dorsal aleurone layer disorders: **a** – L, two cell complexes formed by anticlinal cytokineses shown by horizontal dotted lines and a complex with periclinal cytokineses shown by arrows; **b** – L, discontinuity of the aleurone layer (dotted line); **c** – NH, an elongation growth of aleurone cells below the concave dorsal part (arrow); **d** – NH, an anomalous HP-domain composed of isodiametric cells (arrow) and isolated by aleurone cells (blue) penetrating down the starchy endosperm; **e** – H3, ingrowths of aleurone cells in starchy endosperm (arrows) made by elongation and intrusive growths; **f** – H1, ingrowths of multiplied aleurone cells dividing starchy endosperm into domains (white dots). For **a** – **f** - autofluorescence of tissues.

A key feature of the aleurone layer is its ability to form ingrowths into the starchy endosperm, driven by the elongation and intrusive growth of cells. This results in the isolation of fragments of the starchy endosperm, which can develop differently, such as forming a conical domain of isodiametric HP-subaleurone cells (Fig. 5d). Figure 5e shows the autofluorescent continuity of the aleurone cell walls extending from their outer layer into the starchy endosperm. Periclinal and/or diagonal divisions, combined with elongation and intrusive growth, place typical aleurone cells deep within the starchy endosperm. Observing this phenomenon under normal light microscopy might lead to misleading interpretations, suggesting a possible ‘aleurone’ mutation within the starchy endosperm. The aforementioned penetration of the aleurone layer cells (Fig. 5d) is particularly noticeable in hybrid forms (Fig. 5e, f), resulting in the formation of numerous structural domains within the endosperm (Fig. 5f).

#### 3.2.4. Main morphotypes of starchy endosperm architecture

Three examples of spatial arrangements for the high-protein subaleurone layer and starchy endosperm, containing cylindrical and isodiametric cells, are depicted in Fig. 6. In the Octavia variety (Fig. 6a), the subaleurone layer is similar to the inner part of the endosperm, with occasional cells slightly stained by bromophenol blue. In the Atlas 66 and Nap Hal varieties (Fig. 6b, c), the high-protein subaleurone layer varies in thickness and remains continuous. In all three examples, the distinct long cylindrical cells diminish in the lateral positions (indicated by straight diagonal dashed lines) and gradually shorten towards the isodiametric cells of the endosperm. The isodiametric endosperm can be symmetrically arranged on the sides of the caryopsis (as seen in Atlas 66, Fig. 6b) or asymmetrically (Octavia, Fig. 6a; Nap Hal, Fig. 6c). In Nap Hal, the volume of the isodiametric endosperm is significantly reduced. Near the endosperm cavity, the endosperm is typically composed of isodiametric cells. The shape of cells, derived from periclinal divisions of the aleurone layer surrounding the cavity, suggests their elongation growth towards the dorsal part of the caryopsis. These observations demonstrate intervarietal variability in the development of the HP-subaleurone layer, the spatial relationship between cylindrical and isodiametric endosperm, and the differences in the volume of these tissues.

**Fig. 6.**
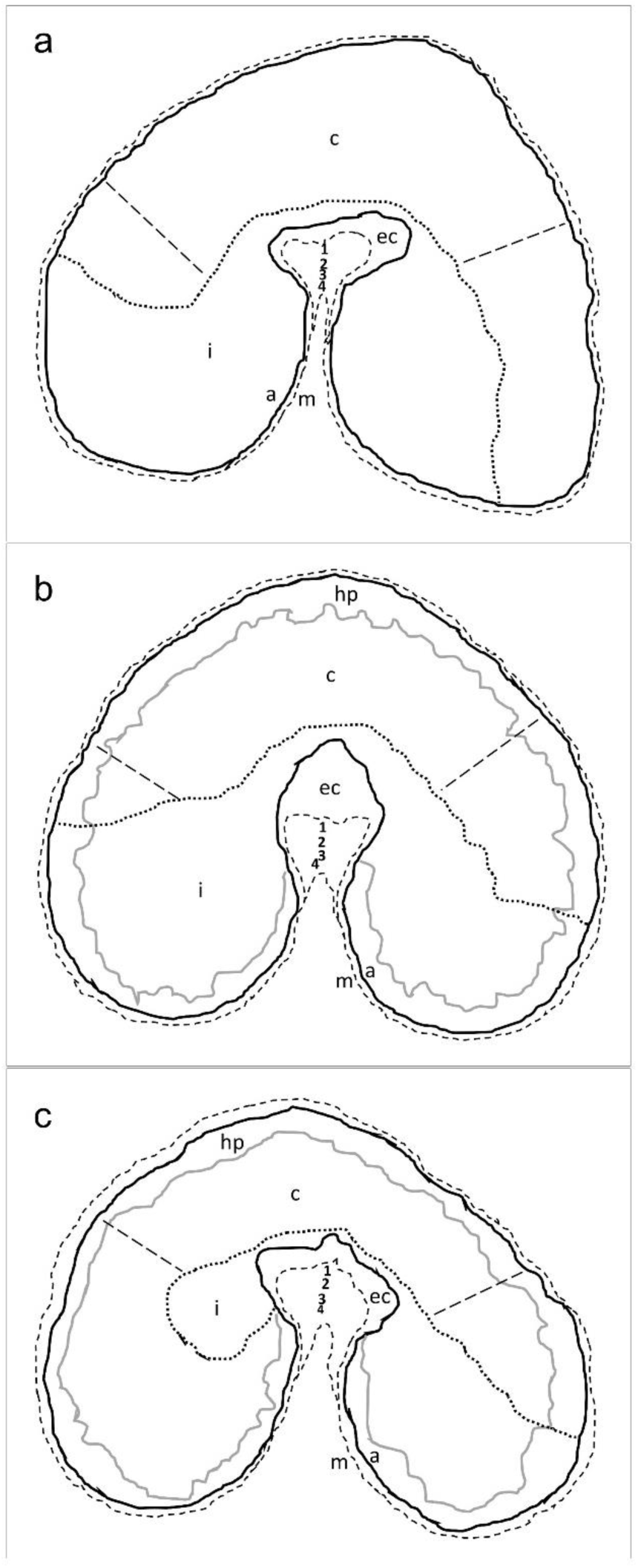
Starchy endosperm architecture based on original microscopic slides: **a** – Octavia (O), **b** – Atlas 66 (A), **c** – Nap Hal (NH). Description of symbols in drawings: a – aleurone layer, c – cylindric endosperm, ec – endosperm cavity, hp – high protein subaleurone layer, i – isodiametric endosperm, m – maternal plant tissues; 1 – nucellar projection, 2 – pigment strand, 3 – xylem bundle, 4 – phloem bundle; two straight dashed lines show area of distinct cylindric endosperm. The drawings are of comparable size.

#### 3.2.5. Characteristics of endosperm cavity structure

In the cross-section of the caryopsis, the endosperm cavity, located within the crease above the vascular system of the grain, can be distinguished (Fig. 7). During development, this cavity becomes filled with the products of apoptotic degradation from the chalazal part of the nucellus. The cavity is separated from the starchy endosperm by a continuous aleurone layer (Fig. 7a-f, arrows), although occasional discontinuities are present (Fig. 7f, arrowheads). Six cavity morphotypes were identified in the common wheat types analysed. While one type tends to dominate quantitatively in a given variety, it co-occurs with other morphotypes, creating a distribution of variability. These morphotypes can be grouped into three developmentally related units, namely: **1** - types A, B, and C (Fig. 7a-c), developing laterally; **2** - types D and E (Fig. 7d, e), developing towards the dorsal part of the caryopsis; and **3** - type F (Fig. 7f), a developmental anomaly related to unit 1 and/or 2. This classification of morphotypes is supported by significant negative correlations between morphotypes from the first and second units (Table 4). Morphotype A does not correlate with any other morphotypes or with the SAC and WEC characters, but it shows intermediate developmental stages leading to types B and C, which justifies its inclusion in the first unit. Morphotype D, characteristic of the Purdue variety, has the highest values for the SAC character and displays significant covariation with it, leading to the highest correlation coefficient between D and SAC. The non-significant correlation coefficients for the anomalous morphotype F are due to its rare, random occurrence, and thus its lack of correlated covariation with other morphotypes.

**Fig. 7.**
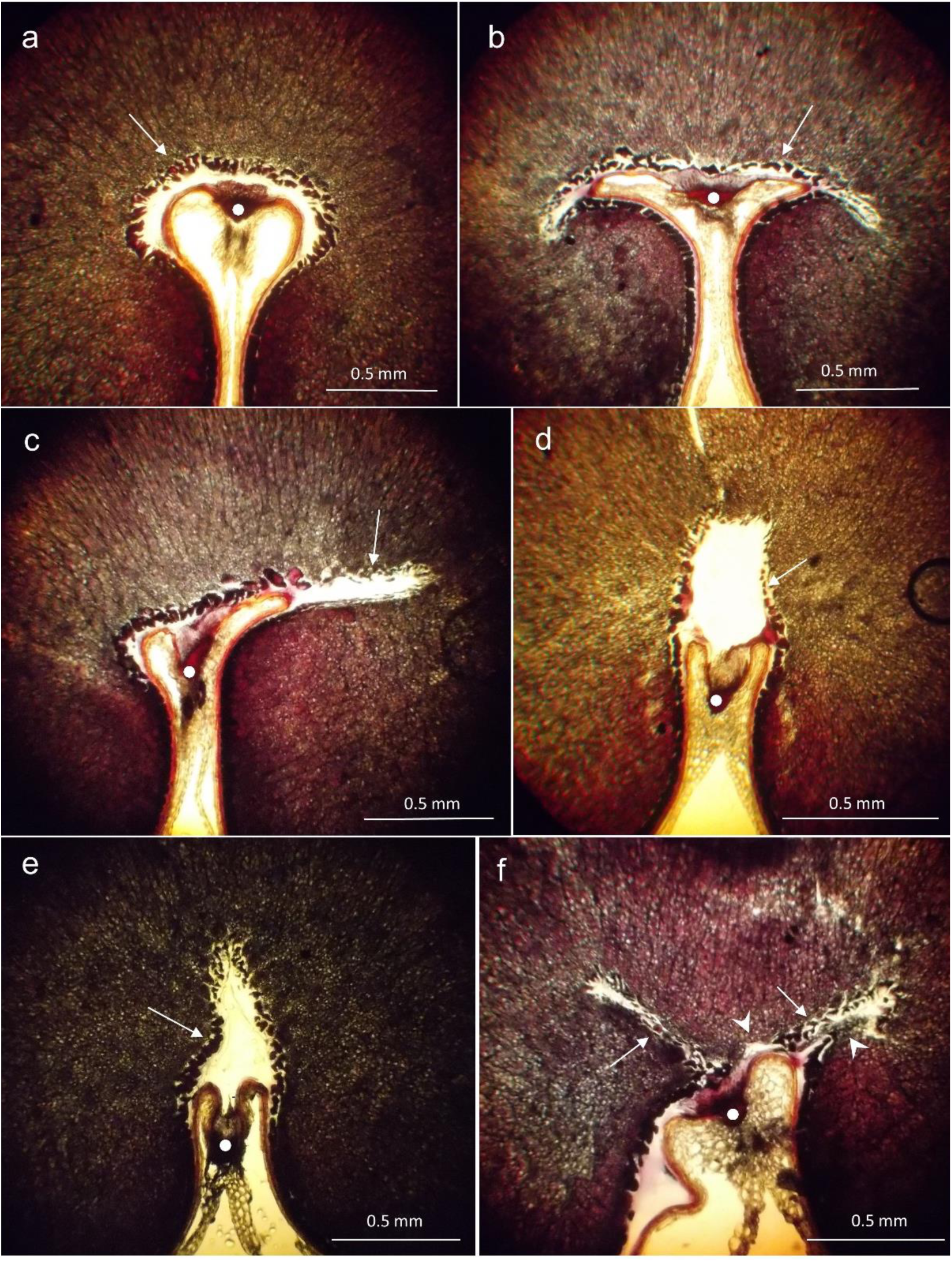
Morphotypes of endosperm cavity in common wheat on cross-sections of caryopsis. **a** – type A (symmetric cavity without lateral arms, H4), **b** – type B (horizontal symmetric cavity with two lateral arms, OP), **c** – type C (horizontal symmetric cavity with one lateral arm, OP), **d** – type D (vertical cavity, it can be also diagonal, L), **e** – type E (vertical narrow cavity often showing intrusive growth of aleurone cells at the top, L), **f** – type F (irregular, anomalous cavity, P). Arrows show the aleurone layer surrounding the cavity and white dots indicate the position of transfer tissues. Arrowheads show discontinuity of the aleurone layer in **f.** For **a** - **f** – cross-sections were stained with bromophenol blue.

**Table 4.**
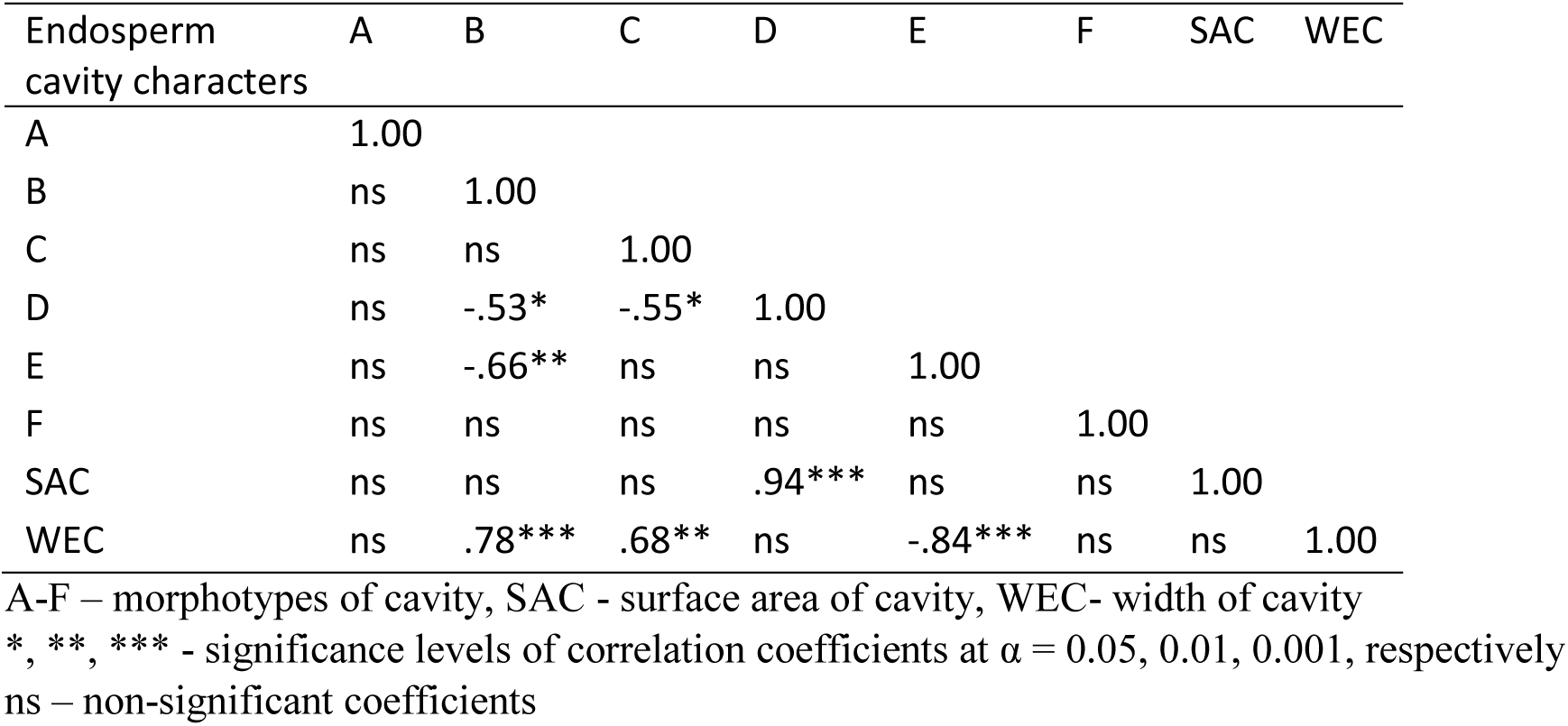
Pearson’s coefficients of correlation between endosperm cavity morphotypes and SAC, WEC characters (*n* = 15 OTUs)

| Endosperm cavity characters | A | B | C | D | E | F | SAC | WEC |
| --- | --- | --- | --- | --- | --- | --- | --- | --- |
| A | 1.00 |  |  |  |  |  |  |  |
| B | ns | 1.00 |  |  |  |  |  |  |
| C | ns | ns | 1.00 |  |  |  |  |  |
| D | ns | -.53* | -.55* | 1.00 |  |  |  |  |
| E | ns | -.66** | ns | ns | 1.00 |  |  |  |
| F | ns | ns | ns | ns | ns | 1.00 |  |  |
| SAC | ns | ns | ns | .94*** | ns | ns | 1.00 |  |
| WEC | ns | .78*** | .68** | ns | -.84*** | ns | ns | 1.00 |

A-F – morphotypes of cavity, SAC - surface area of cavity, WEC- width of cavity
\*, \*\*, \*\*\* - significance levels of correlation coefficients at $\alpha = 0.05, 0.01, 0.001$ , respectively
ns – non-significant coefficients

#### 3.2.6. Developmental events in the area of the endosperm cavity

The endosperm cavity is formed between the modified aleurone layer (MAL) in the caryopsis crease region and the tissues of the ovule chalaza, which have transformed into the nucellar projection (**np**). The pigment strand (**ps**) and the vascular bundles of the xylem and phloem, which are geometric extensions of the chalaza, are components of the nucellus, an organ of the maternal plant. The MAL typically consists of a single layer of thick-walled cells, often exhibiting intrusive wall growth towards the starchy endosperm (Fig. 8a). In some areas, MAL can be multi-layered, with two or more cell layers. The cavity interior is sometimes filled with apoptotic remnants of the nucellus, which rarely retain a cellular structure (Fig. 8b).The multi-layering of MAL, along with the contacts between aleurone cells and adjacent starch cells (Fig. 8c, g), indicates that MAL is meristemically active, contributing daughter cells to the starchy endosperm through periclinal divisions. A comparison with the dorsal aleurone layer (DAL) shows that DAL is even more meristematically active. MAL cells are frequently polyploidised (Fig. 8g). The cavity is not empty but is always filled with nucellar remnants, which are typically amorphous (Fig. 8d-f).The intensive elongation and intrusive growth of MAL and DAL cells lead to connections between these layers, creating barriers that facilitate the formation of endosperm domains (Fig. 8e). Discontinuities in the MAL layer are common (Fig. 8f-i), allowing direct contact between the cavity’s contents and the starch of the endosperm. These discontinuities enable the penetration of metabolites and/or the synthesis of starch grains within the cavity (Fig. 8h, i). This morphotype is found with varying frequencies in different cultivars, occurring at 14% in the Purdue cultivar, 5% in Lancota, and 1% in Nap Hal.

**Fig. 8.**
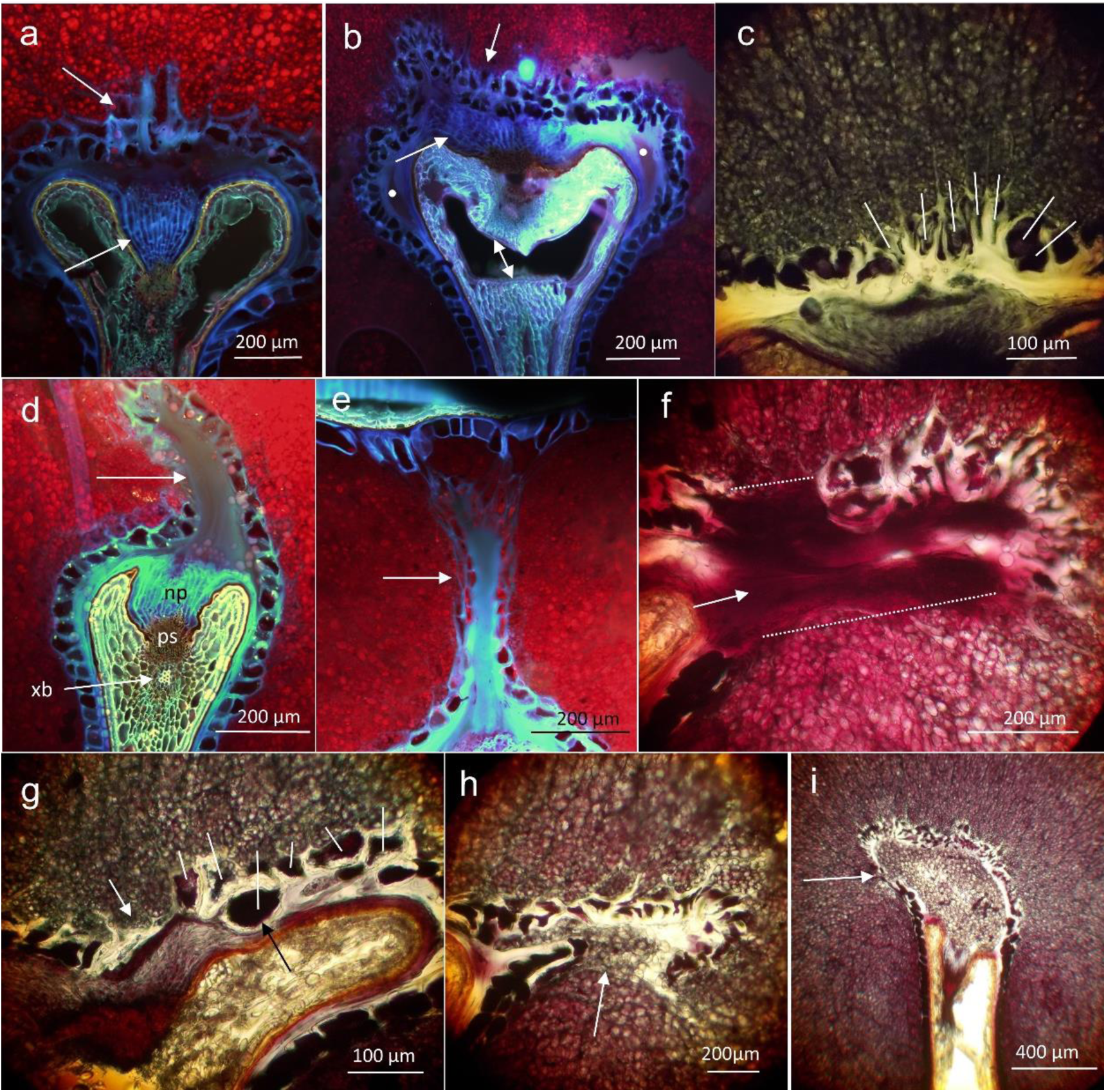
Developmental events in the endosperm cavity. **a** – CI (upper arrow shows the thick-walled modified aleurone layer (MAL) with wall ingrowths into starchy endosperm, lower arrow shows nucellar projection);, **b** – H8 (upper arrow shows the thick-walled, multi-layered MAL, lower arrow shows a fragment of nucellar tissue in the cavity, two white dots mark amorphous parts and two-headed arrow for an artifact of teared nucellar projection); **c** – H3 (white lines indicate developmental complexes of MAL cells and sister starch cells formed after tangential divisions); **d** – H8 (upper arrow shows an amorphous content of the cavity, and below symbols for: np – nucellar projection, ps – pigment strand, xb – xylem bundle); **e** –NH (arrow shows an amorphous blue content of the cavity and connection of MAL with the dorsal aleurone layer); **f** – P (arrow shows an amorphous red content of the cavity and dotted lines MAL discontinuity); **g** – OP (white lines indicate developmental complexes of MAL cells and sister starch cells formed after tangential divisions, white arrow for MAL discontinuity and black arrow shows a polyploid MAL cell); **h**- P (arrow marks MAL discontinuity and penetration of starch in the cavity); **i** – P (arrow indicates MAL discontinuity and the cavity filled by starch grains). Autofluorescence of tissues in **a**, **b**, **d**, **e**, and bromophenol blue staining in others.

#### 3.2.7. Distribution of endosperm cavity characters in nmMDS ordination space

In the ordination space, three structural characteristics of the cavity are distinguished (Fig. 9): 1. SAC (the most distant from the others). 2. WEC. 3. All cavity morphotypes that have close distances between them. SAC is the most synthetic characteristic of the cavity, influenced by numerous determinants of its morphogenesis, while WEC reflects the lateral growth of the MAL and cavity, with fewer determinants compared to SAC. The mutual positioning of the cavity morphotypes in the minimum spanning tree (MST) illustrates their morphogenetic similarities, as shown in Fig. 7, with morphotype A centrally located, indicating its role as the initial form for the development of other types.

**Fig. 9.**
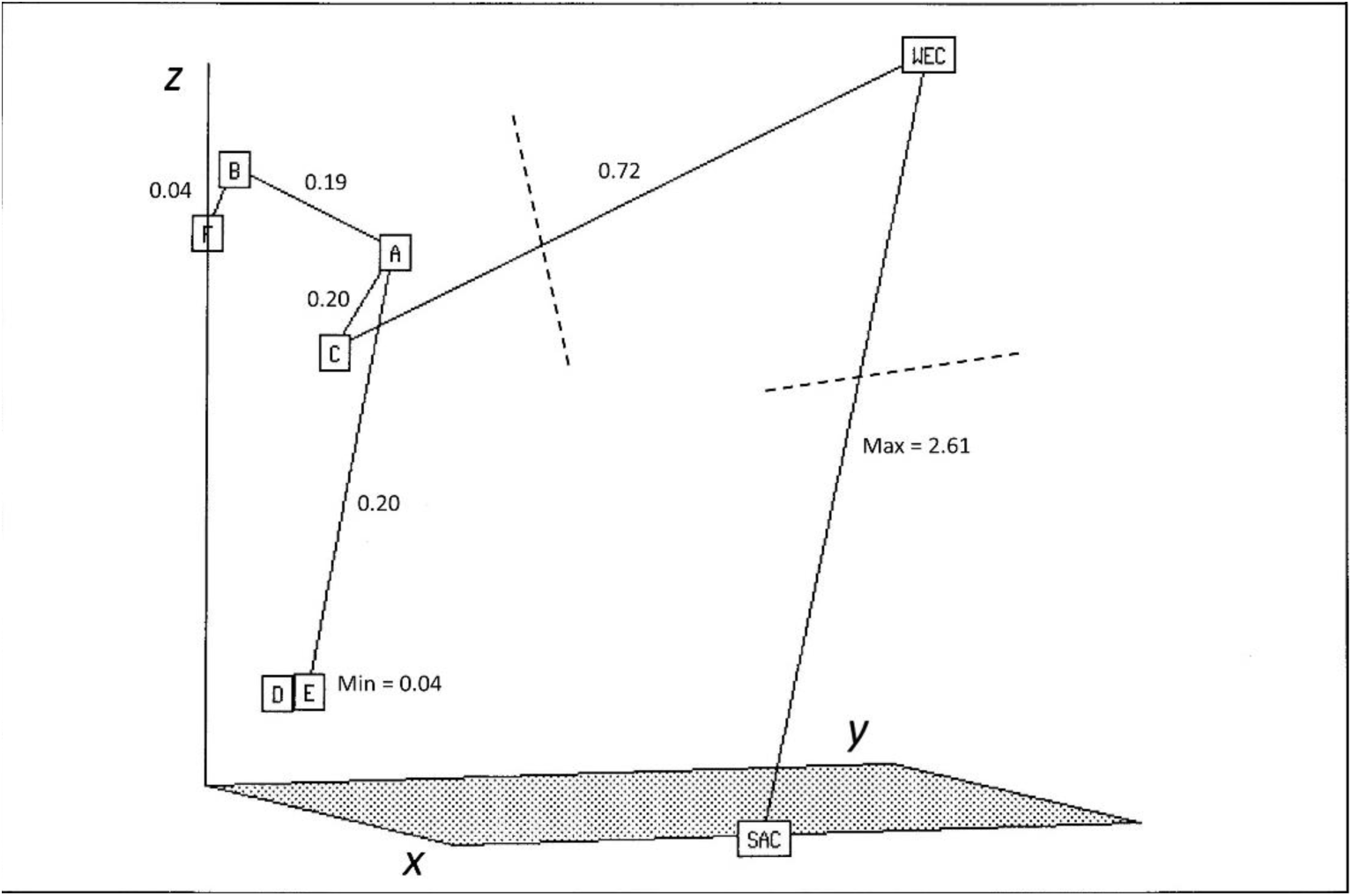
Minimum spanning tree of eight endosperm cavity characters in an ordination space defined by *x*-, *y*-, and *z*-axes value, and created by the application of Kruskal’s non-metric multidimensional scaling method [27](Rohlf 1994). Characters were described by 15 arithmetic averages of common wheat breeds. In the tree minimum and maximum and other distances are presented. Labels for characters in M&M and Fig. 7.

#### 3.2.8. Distribution of common wheat OTUs described by the frequency of endosperm cavity morphotypes in nmMDS ordination space

In the MST diagram, two types of wheat forms are distinguished based on the frequency distribution of endosperm cavity morphotypes, excluding SAC and WEC characters: 1. A type with a dominant single cavity morphotype, corresponding to a leptokurtic distribution, and 2. A type with co-dominant two or more cavity morphotypes, corresponding to a platykurtic distribution. At the center of the MST diagram (Fig. 10) are wheats of type 1, including A, P, H2, and the H3 hybrid, which exhibits four cavity morphotypes of similar frequency. Such variation is observed in both varieties and hybrids. On the periphery of the diagram are wheats of both types 1 and 2; however, OTUs with similar cavity morphotypes are often directly connected, such as CI (E) and H7 (AE) or H6-O (AC) and H4 (ABC). Morphotype D is dominant (50%) in the Lancota (L) variety and H5 hybrid. However, a great distance is present between L and H5 in the MST tree (Fig. 10), resulting from the difference between the distribution of morphotype frequencies, and especially in relation to co-dominant morphotypes, C (23%) and A (32%), respectively.

**Fig. 10.**
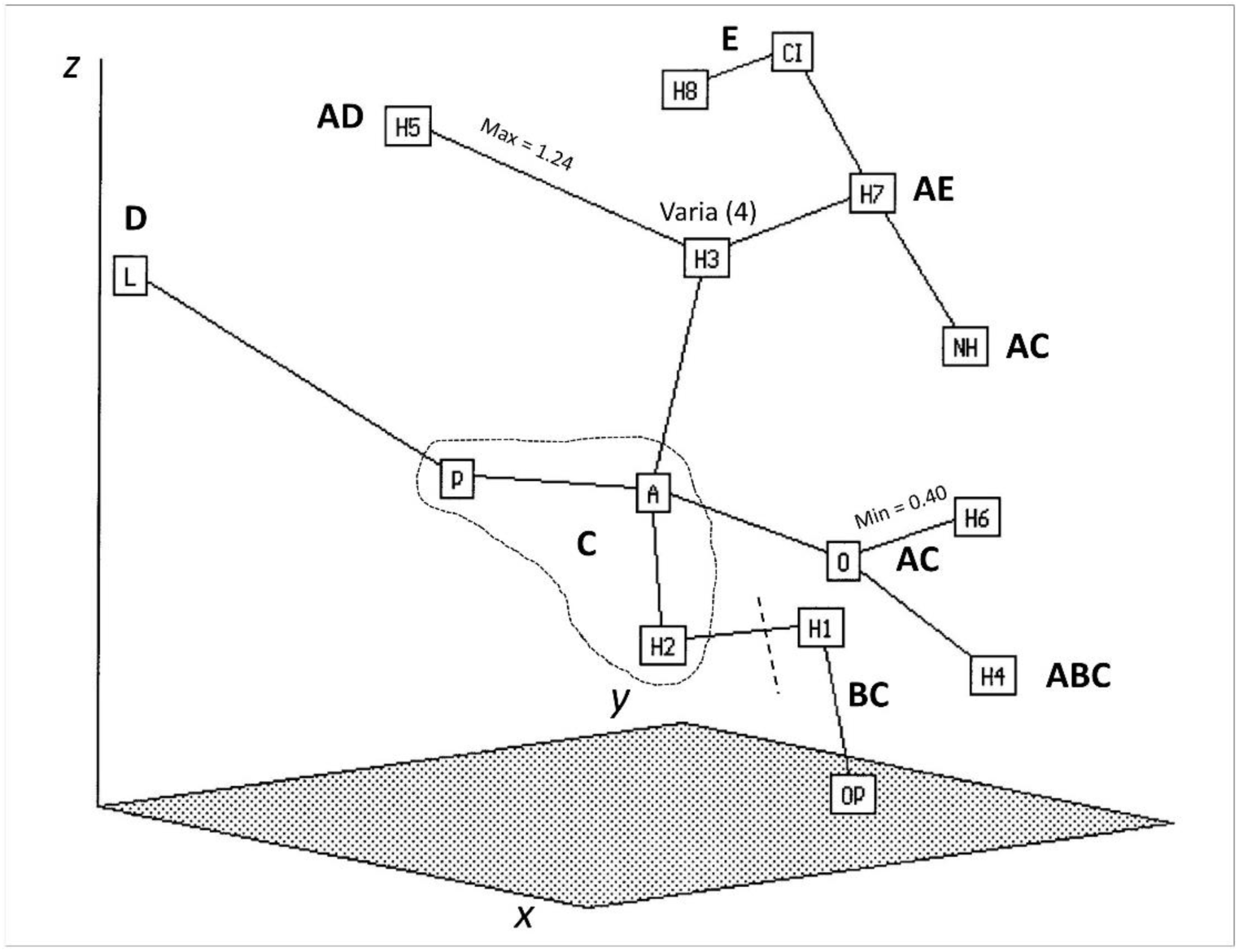
Minimum spanning tree of 15 OTUs of common wheat described by the frequency of six morphotypes of endosperm cavity in an ordination space defined by ***x***-, ***y***-, and ***z***-axes, and created by the application of Kruskal’s non-metric multidimensional scaling method [27](Rohlf 1994). SAC and WEC characters were omitted here. In the tree, minimum and maximum distances are presented. Most frequent morphotypes (A, B, C, D, E) in OTUs are written in bold nearby labels for OTUs. Labels for OTUs are in M&M.

## 4. Discussion

### 4.1. Interrelations of caryopsis characters and OTUs

Vitreousness is related to the compactness of the endosperm, which depends on protein content, the gliadin/glutenin ratio, and protein-starch adhesion [5,29]. This relationship was confirmed by positive correlations between SG and MV. The correlations of SG with the thickness of the high-protein subaleurone layer (HP-subaleurone) were positive in some OTUs (e.g., CI), negative in others (NH, P, H2, H4, H8), or insignificant in the rest, indicating that, besides protein content, other structural components of the caryopsis, such as the size of the endosperm cavity, significantly influence SG. The W-SAC correlation proved that larger mature caryopses tend to have larger endosperm cavities, a developmental trend also noted by [21]. The structural variability of the caryopsis across OTUs was confirmed using PCA, which revealed significant covariation in trait groups. Principal components distinguished information about ‘caryopsis size’ from ‘caryopsis quality’, a separation similar to that found in the Opolska cultivar from Poland [6]. The distribution of characters in the ordination space, based on average taxonomic distances, separated components such as SG, MV, and SAC from quality related traits like TA, TDS, and TLS. In the ordination space, two groups of OTUs - described as HP-subaleurone and LP-subaleurone - were identified, with an intermediate cluster of three cultivars. Protein analyses from the University of Nebraska, Lincoln (VA Johnson, pers. comm.) showed that Lancota, with the highest protein content (13.1%) in this group, was closest to the HP-subaleurone group. Historically and currently, the HP-subaleurone varieties Nap Hal, Atlas 66, and Lancota have been identified as genetically diverse sources of high protein content in the caryopsis [1,3,4,30].

### 4.2. Details of caryopsis microstructure

In the developing wheat seed,the chalazal region forms before the endosperm in the embryo sac. The nucellus undergoes significant apoptotic degradation [20], while its chalaza transforms into a nucellar projection (**np**), showing varied morphogenesis and vertical or horizontal cell orientation [12]. Thick-walled nucellar projection cells distinguish this structure and assign it a special morphogenetic role [8]. Depending on the **np** shape, two extreme endosperm cavity morphotypes - *planar* or *gothic* - influence cylindrical starchy endosperm architecture. The dorsal aleurone endosperm shows more meristematic activity and ordered cytokineses, mainly tangential, compared to the less organised ventral part. In the dorsal part, tangential cytokineses are not frequent, when the daughter cells undergo intensive elongation and intrusive growth, with the penetration of sharp, narrow cell ends after diagonal division into intercellular spaces. Numerous examples of intrusive growth in the plants’ endosperm were observed at the tips of haustorial protrusions [31]. In the endosperm of *Thinopyrum distichum*, the dorsal and ventral cylindric cell complexes showing intrusive growth were very distinct, with small morphogenetic differences between them [32]. In wheat cv. Crousty, intrusive growth in the starchy endosperm occurred after diagonal cytokineses of cylindrical cells [29]. In interspecific amphiploids of *Avena*, intrusive growth was demonstrated by cells of the aleurone layer, and free spaces in the endosperm additionally enabled it. Such growth of aleurone cells after their divisions caused them to penetrate the starchy endosperm, which can be incorrectly interpreted as a mutation [15]. However, the *globby1-1* mutation of this type discovered in maize is real [33]. The allocation of energy resources to intensive growth instead of cytokinesis may result in the morphogenesis of a long cell, with an unchanged phenotype from the initial aleurone layer, penetrating deep into the starchy endosperm. Examples of such morphogenesis, both for starch cells (rarely) and aleurone cells (often), were documented in this article. Such single events were more frequently noted in developmentally unstable hybrid forms than in stable varieties and concerned larger endosperm domains that were developmentally different [15]. As shown (Fig. 5d-f), a special role in the formation of endosperm domains is played by diagonal divisions and intrusive growth of aleurone cells, which is enhanced by apoptotic-like elimination of nuclei from the syncytial embryo sac [14–16]. It has been previously shown that the elimination of nuclei at this stage causes disturbances in endosperm development in *Triticale* [34]. Anomalous endosperm development in rye, wheat, and *Triticale* was also explained by premature amylase activity [35]. Numerous barley mutants with anomalous endosperm and acellular spaces were characterised by callus-like differentiation of globular groups of cells [36], similar to the *globby1-1* mutation in *Zea mays* [33]. Morphogenetic choices between directional or chaotic cytokineses may concern entire individual caryopses with isodiametric vs cylindrical endosperm in *Avena* [13]. However, most often, this selection is balanced - see ‘morphotypes of endosperm architecture’ (Fig. 6). In starchy endosperm, a subaleurone layer of cells is distinguished, which forms after the last tangential divisions of the aleurone layer. The cells of this layer have a different storage potential compared to the central endosperm. [37] showed that this layer has a higher protein content and fewer starch grains. It has also been proven that small vascular bundles in the caryopsis correlate positively with a thicker HP-subaleurone layer [38]. This could represent a state of starvation in the caryopsis, leading to decreased starch synthesis and a shift in proportion favouring protein. See the differences in grain structure between xeromorphic steppe varieties and hygromorphic varieties of Western Europe [39], which result from climate differences (precipitation, temperature) and are also dependent on the length of the vegetation period [40 after Flaksberger]. This was confirmed by drought stress studies, showing a deficiency of B-type starch grains in the subaleurone layer [41]. The development of the HP- subaleurone layer is critical for wheat grain quality and varies across the OTUs studied. However, its absence around the endosperm cavity have been demonstrated as a stable characteristic. It should be added that a HP-subaleurone part can be single- or multi-layered, in terms of cells. High-protein types like Atlas 66 and Nap Hal, used in breeding programs [3], have a distinct continuous HP-subaleurone layer, while varieties like Octavia lack it (see Fig. 4). In high-protein varieties, a clear gradient of protein to starch exists from the periphery to the center, unlike in varieties where starch synthesis is uniform throughout the endosperm (cv. Octavia). The variability in HP-subaleurone wheat varieties is considered typical in other cereals and concerns subaleurone, cylindrical and central isodiametric endosperm compartments. [42] suggest that increasing the protein content from transfer tissues towards the aleurone layer is associated with the expression of gluten proteins encoding genes in the subaleurone layer.

### 4.3. Characteristics of endosperm cavity

It has been demonstrated that the breeding unit of common wheat can be characterised by the dominant type of endosperm cavity in the distribution of variation of cavity type frequency. While the entire distribution provides a complete picture, it also introduces complexity. The distributions studied presented a larger cumulation of variation (leptokurtosis) or a smaller cumulation (platykurtosis). The shape of the cavity is influenced by the shape of the caryopsis [43], which itself depends on the shape of the floral cavity formed by the surrounding glumellae [17,44]. The floral cavity also affects the morphotypes of wheat lodicules that develop between the caryopsis and glumellae [18], indicating a correlated development between caryopsis and lodicule morphotypes. These interrelations within the flower cavity align with Davenport’s findings on the developmental organisation of flowers in *Ranunculus ficaria* [45]. It was proven that flower organs that are closer to each other in the geometric sense are more highly correlated. It would also mean that any development of structures/tissues (lodicules, caryopsis, endosperm cavity, starchy endosperm) within a wheat lemma-palea cavity can be correlated. One such relationship could be the influence of the of the caryopsis shape, whether a domed or flat, on the ventral-dorsal architecture of the starchy endosperm. The endosperm cavity, part of the apoplast in the caryopsis, contains high concentrations of sucrose and oligosaccharides [5,21,22], which supports *in situ* starch synthesis. Assimilates for starch production are supplied via the transfer system, including the nucellar projection. However, MAL discontinuities and the direct contact of starch grains within the cavity and the cellular endosperm suggest that both regions form a single ‘starchy’ compartment. The endosperm cavity also contains arabinoxylans, synthesised in the nucellar epidermis, which regulate caryopsis hydration [46]. In most common wheat breeds, the cavity is predominantly astructural (Fig. 8d-f), though it sometimes contains cells or tissue fragments (Fig. 8b). This characteristic is also shared by the cavity in the amphiploid *Triticum timopheevii/Aegilops umbellulata* [47].

## 5. Concluding remarks

The analysis of the caryopsis characters revealed that specific gravity (SG) is influenced by endosperm cohesion (MV character) and endosperm cavity volume. The studied OTUs differ in the loadings of characters in the subsequent principal components; however, components I (caryopsis weight), II (caryopsis quality), IV (mealyness vs. vitreousness), and V (aleurone layer) provide information on the variation of these characters in most OTUs. Information on SG variation is distributed across several components. In the ordination space, caryopsis quality characters are distinguished from others. In the minimum spanning tree, OTUs are grouped into HP-subaleurone and LP-subaleurone clusters. The architecture of the endosperm can be depicted by the arrangement patterns of the HP-subaleurone layer and the cylindrical and isodiametric starchy cells. The dorsal part of the starchy endosperm is developmentally shaped by the *planar* or *gothic* endosperm cavity, with cell elongation and intrusive growth, particularly in the aleurone layer, playing a significant role in forming different endosperm domains. The endosperm cavity, which develops above the caryopsis transfer system, also has a key morphogenetic function. Six cavity morphotypes have been identified, developing according to *planar* and *gothic* patterns. In addition, acellular domains were discovered in the starchy endosperm, and a rare occurrence of starch grain synthesis was observed in the endosperm cavity, both events being cultivar-specific, and more frequent in the Purdue cultivar.

## Acknowledgements

The authors thank Prof. P. Stephen Baenziger, Lincoln, University of Nebraska, for his kind help in identifying the origin of some wheats.

## CRediT authorship contribution statement

**RK:** Scientific conception, Investigation, Data curation, Numerical analysis, Writing the manuscript. **PT:** Fluorescence microscopy data analysis and respective Figs, Read, corrected and supplemented the manuscript.

